# Adaptation to space conditions of novel bacterial species isolated from the International Space Station revealed by functional gene annotations and comparative genome analysis

**DOI:** 10.1101/2023.09.28.559980

**Authors:** Lukasz M. Szydlowski, Alper A. Bulbul, Anna C. Simpson, Deniz E. Kaya, Nitin K. Singh, Ugur O. Sezerman, Paweł P. Łabaj, Tomasz Kosciolek, Kasthuri J. Venkateswaran

## Abstract

**Background:** The extreme environment of the International Space Station (ISS) puts selective pressure on microorganisms unintentionally introduced during its 20+ years of service as a low-orbit science platform and human habitat. Such pressure leads to the development of new features not found in the Earth-bound relatives, which enable them to adapt to unfavorable conditions.

**Results:** In this study, we generated the functional annotation of the genomes of five newly identified species of Gram-positive bacteria, four of which are non-spore-forming and one spore-forming, all isolated from the ISS. Using a deep-learning based tool - deepFRI - we were able to functionally annotate close to 100% of protein-coding genes in all studied species, overcoming other annotation tools. Our comparative genomic analysis highlights common characteristics across all five species and specific genetic traits that appear unique to these ISS microorganisms. Proteome analysis mirrored these genomic patterns, revealing similar traits. The collective annotations suggest adaptations to life in space, including the management of hypoosmotic stress related to microgravity via mechanosensitive channel proteins, increased DNA repair activity to counteract heightened radiation exposure, and the presence of mobile genetic elements enhancing metabolism. In addition, our findings suggest the evolution of certain genetic traits indicative of potential pathogenic capabilities, such as small molecule and peptide synthesis and ATP-dependent transporters. These traits, exclusive to the ISS microorganisms, further substantiate previous reports explaining why microbes exposed to space conditions demonstrate enhanced antibiotic resistance and pathogenicity.

**Conclusion:** Our findings indicate that the microorganisms isolated from ISS we studied have adapted to life in space. Evidence such as mechanosensitive channel proteins, increased DNA repair activity, as well as metallopeptidases and novel S-layer oxidoreductases suggest a convergent adaptation among these diverse microorganisms, potentially complementing one another within the context of the microbiome. The common genes that facilitate adaptation to the ISS environment may enable bioproduction of essential biomolecules need during future space missions, or serve as potential drug targets, if these microorganisms pose health risks.

## 1 Introduction

The environment of outer space, characterized by intense radiation and high vacuum, presents harsh conditions for any form of life. Despite this, microorganisms have demonstrated a remarkable adaptability to these extreme space environments. Organisms known as extremophiles, such as bacteria (e.g. *Bacillus, Deinococcus* species) [1, 2], fungi [3], tardigrades [4], bdelloid rotifers [5], and many others, are well-documented for their ability to survive in space conditions. As such, it is virtually impossible to prevent the transfer of microbial life to exploratory space vessels or planets[6, 7]. In addition to microorganisms potentially hitching a ride with robotic components, the microbiomes of humans also travel to space. This not only includes the microbiomes of space crew members, but also free-living microorganisms associated with cargo, science instruments, and test animals, including potential pathogens which should be considered for their potential to adapt to space conditions [8–13].

The exposure of microbes to outer space, or simulated versions of these conditions, has been extensively studied to test survival capabilities and assess the panspermia hypothesis. Over several decades, various microbial samples have been exposed to space conditions via balloons, rockets, and spacecraft as part of the pioneering experiments in astrobiology (reviewed in [14]). For example, after 18 months of exposure, *Bacillus pumilus* SAFR-032, a bacterium isolated from the Jet Propulsion Laboratory’s Spacecraft Assembly Facility (JPL-SAF), exhibited 10-40% survivability when UV was masked [1]. However, when these SAFR-032 spores were exposed to UV light (above 110 nm) and vacuum in space for 18 months, except for a few spores, there was a roughly 7-log reduction in viability. Similarly, dried *Deinococcus* cell pellets, particularly those 500 *µ*m in thickness, remained viable after three years of space exposure due to a shadowing effect. Moreover, *D. radiodurans* after being subjected to 1-year exposure to outer space during Tanpopo mission[15], showed a collection of molecular alterations e.g. DNA damage and oxidative stress response, accumulation of metal transporters etc. In a recent study, a survival of *D. radiodurans* exposed to near space conditions was correlated with a growth condition prior to the exposure[16]. A comparative proteomics analysis of *B. pumilus* SAFR-032 revealed that proteins conferring resistance traits, such as superoxide dismutase, were present in higher concentrations the vegetative cells derived from spores exposed to space compared to ground controls. Furthermore, the first-generation spores of *B. pumilus* SAFR-032 resulting from space-exposed samples demonstrated increased UVC resistance (around 4,000 J/m^2^) compared to their ground control counterparts[1].

The International Space Station (ISS) is a closed environment that houses a diverse range of microorganisms, including potential pathogens [17]. As we prepare for long-term space explorations, understanding the molecular mechanisms that allow microorganisms to survive and adapt during spaceflight is becoming increasingly important [10, 15]. Monitoring the microbial population onboard the ISS is crucial for safeguarding astronaut health and preventing contamination of both the spacecraft and its equipment [7]. Whole-genome sequencing (WGS) and phenotypic characterization of any novel microbial species discovered on the ISS can help identify potential pathogens and comprehend their impact on the closed habitat and crew health [7]. However, to accurately identify potential microbial threats, it is essential to maintain a current and comprehensive database of microbial genomes, detailing both their species identities and phenotypic traits. This database, continuously updated through WGS of pure cultures, is crucial for gathering genetic data and assisting in the identification of microbial threats via shotgun metagenomic sequencing. Moreover, the active initiatives to characterize the cultivable microbiome of previously not sequenced space habitats contribute to decreasing the unidentified microbial ‘dark matter’ in metagenomic sequencing outcomes.

Research conducted in space or using ground-based simulators has shown that microgravity conditions can alter various biological processes, including growth, morphology, gene expression, virulence, drug resistance, biofilm formation, and secondary metabolism [8–10, 12, 18]. Studies focusing on the microbiome have reported an increase in antibiotic resistance markers in astronauts’ saliva [19], although these changes were temporary and reversed upon returning to Earth. Other research has indicated that microbial isolates from space environments are differentiating themselves from their Earth-bound counterparts, showing signs of adaptation to their new environment [17, 20–23]. Interestingly, under spaceflight conditions, one species, *Bacillus safensis* JPL-MERTA-8-2, exhibited a 60% improvement in growth compared to its Earth-based growth [24].

This study focused on the genomic peculiarities of novel species isolated from the ISS and compared WGS of the closest relatives found in Earth environments. A thorough analysis was conducted on genes associated with DNA repair, radiation resistance, microgravity adaptation, stress responses, and metabolic changes (Fig. 1). These genes were compared to their terrestrial counterparts, and potential variations induced by space conditions were observed. The protein-coding sequences from each species pair were compared using a set of similarity metrics, unearthing potentially novel genes and clusters that might have been evolved in space. Furthermore, mutations within these sequences were rigorously analyzed, and their protein structures were predicted using *in silico* models. This approach illuminates the genetic adaptations undertaken in stringent conditions of the space environment. In essence, by comparing space and Earth genomes and identifying mutation patterns, a clearer insight into genetic evolution influenced by space conditions has been obtained.

**Fig. 1.**
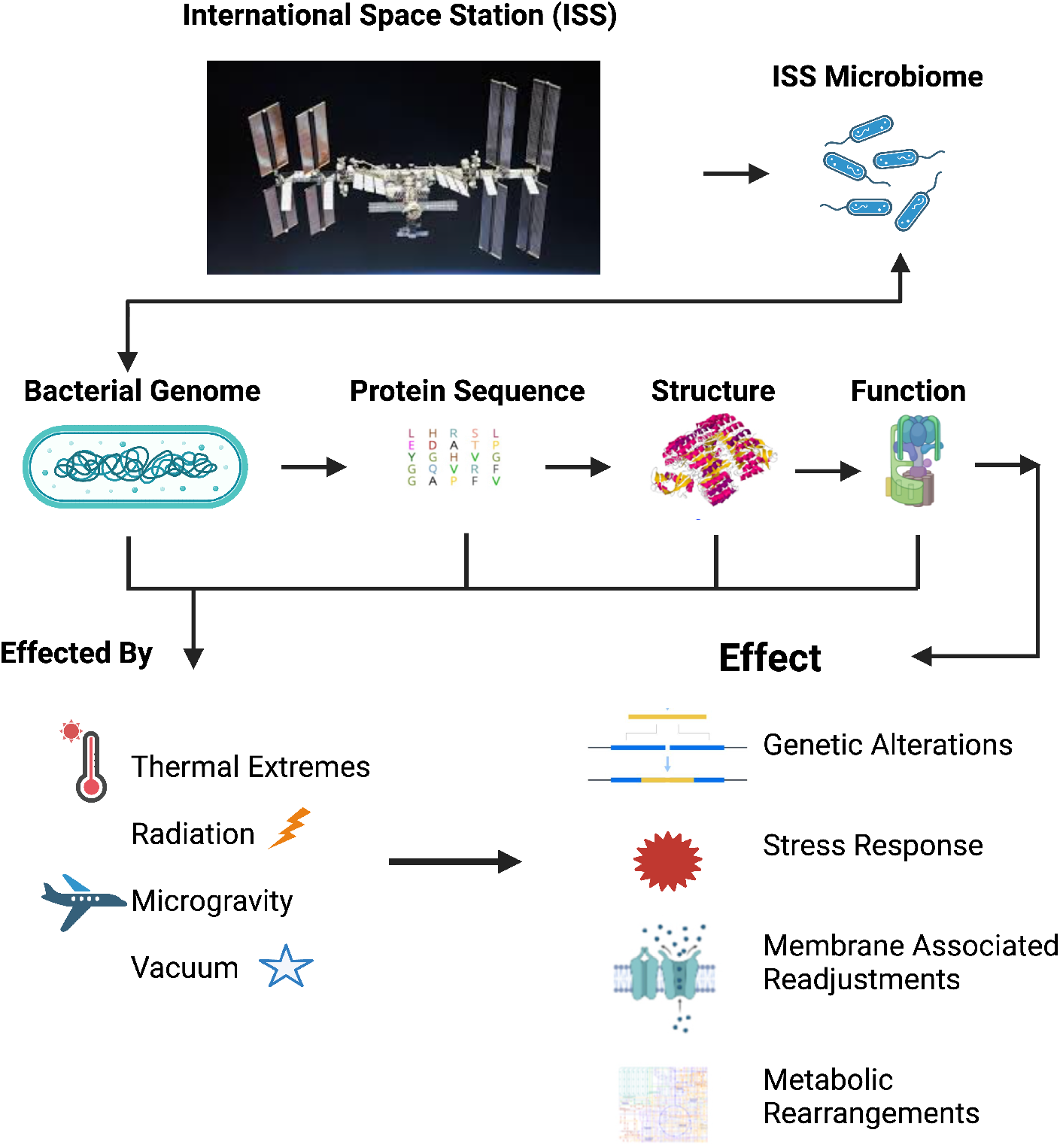
Overview of the project.

## 2 Results

### 2.1 Comparison of different annotations and between the isolates and within pangenome context

Five novel species of Gram-positive bacteria that were isolated from the ISS were analyzed. Their generic features and the results of other molecular analyses are presented in Table 1 and Supplementary Figure S1. These bacteria were obtained from various flights, locations, and time periods, and are associated with different phylogenetic groups. The strain F6 3S P 1C, which belongs to the *Paenibacillus* genus, has been identified as a spore-former, while the other four species were identified as non-spore-forming Actinobacteria. Through ANI and AAI analysis, we established closest Earth relatives (Table 1 and Supplementary Table S1). Additionally, we performed synteny analysis [25] using all top ANI hits for each of the five organisms (Supplementary Table S1), but yielded no results (data not shown), thus indicating all ISS isolates are distinct species.

**Table 1.**
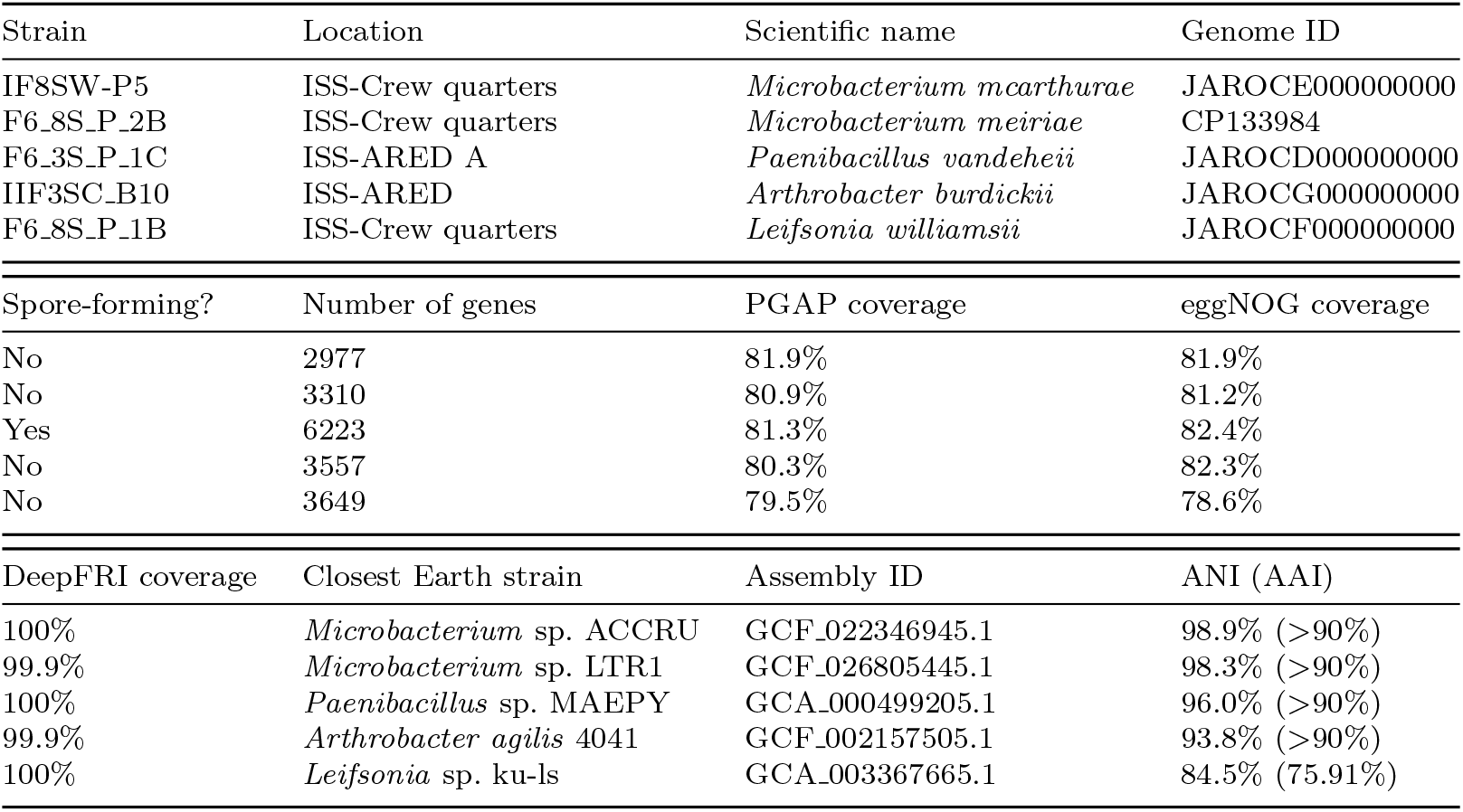
General features of the isolates used in this study. **ANI** - average nucleotide identity.

#### 2.1.1 Annotations common for space and Earth species (closest relative, see Supplementary Figure S1 for more comparisons)

When the annotated genomes were compared with several methods (PGAP, eggNOG), DeepFRI analysis showed highest annotation coverage and provided a confidence scale (based on the score value), to filter medium quality (MQ) and high-quality (HQ) annotations and the results are shown in Figure 2 (see Methods for details). In total, the genomes of the novel species encompassed 1290 and 1184 shared MQ and HQ annotations (1778 all annotations, Fig. 2ab). The majority of shared annotations were related to metabolism, followed by cellular processing and signaling and information storage and processing (Fig. 2c; all common annotations (GO terms) are detailed in Supplementary Table S12 in the Supplementary material).

**Fig. 2.**
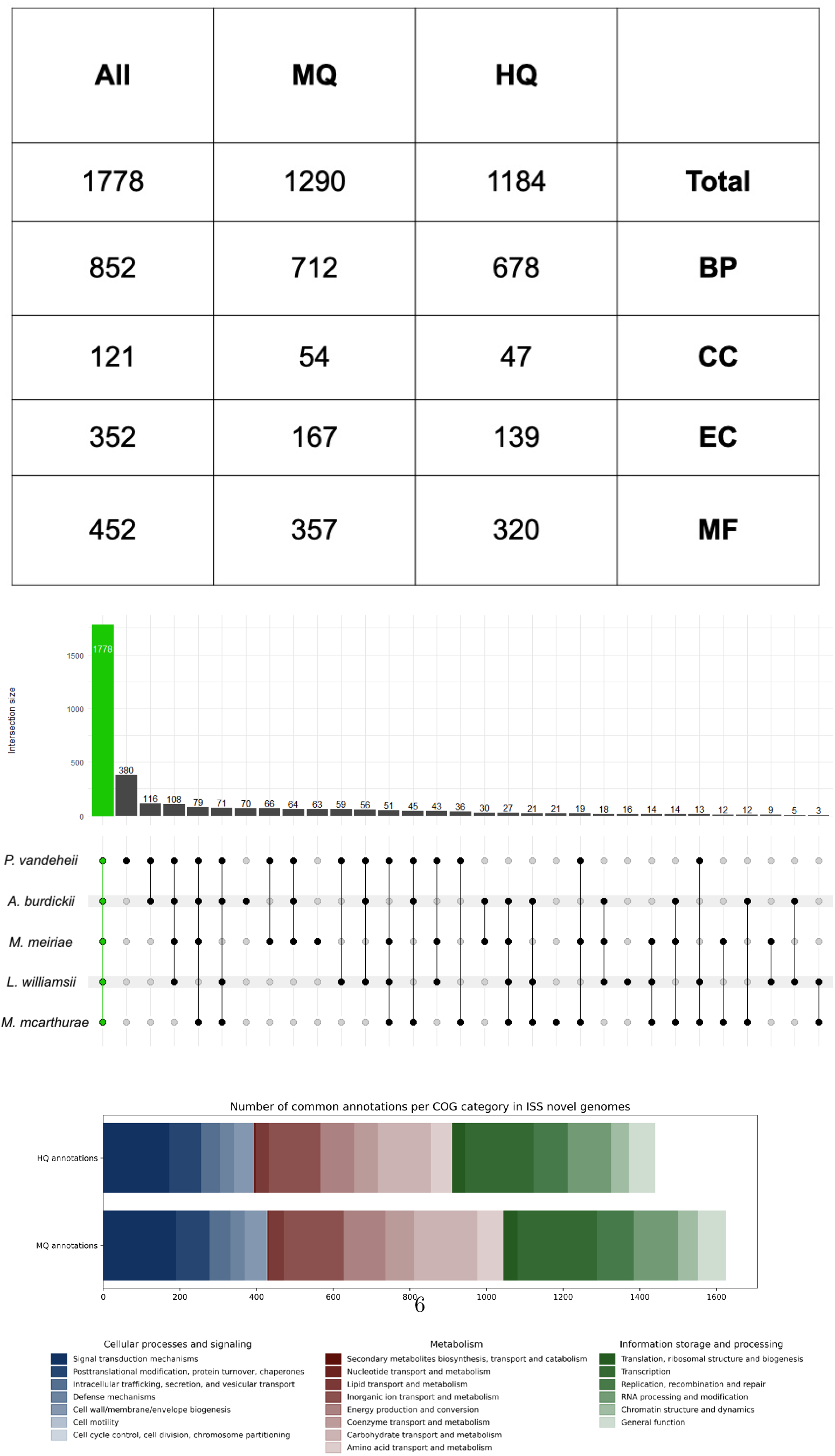
Overview of the annotations common in all ISS species. A) Number of unique, common annotations found in all species (BP - biological process, CC - cell component, EC - enzyme commission, MF - molecular function, MQ - medium quality, HQ - high quality) B) Upset plot showing common annotations (derived from DeepFRI) with different groups of interests C) Clustering all isolates’ common annotations in the form of clusters of orthologous genes (COG) categories. (See Methods for more details.

#### 2.1.2 Annotations unique to ISS bacterial species

All curated National Center for Biotechnology Information (NCBI) genomes were used to find the closest terrestrial relatives for the novel ISS bacterial species and the genome that exhibited the highest average nucleotide identity (ANI) value was further compared with genomes of ISS strains to characterize unique annotation. When compared with their closest terrestrial relatives all ISS species have different numbers of unique annotations (Fig. 3a), which corresponds to their ANI (the higher the ANI, the fewer unique annotations). Additionally, two-dimensional Conserved Domain (CD)-hit clustering was performed between each isolate and its closest relative, using a 40% sequence similarity threshold. Thus, for each isolate a subgroup of unique genes (with no counterparts in their closest relative) was found (Supplementary Tables S2-S11). A comparison between these genes revealed the presence of 74 and 49 MQ and HQ common annotations, respectively (309 total annotations, Fig. 3b, Supplementary Table S13).

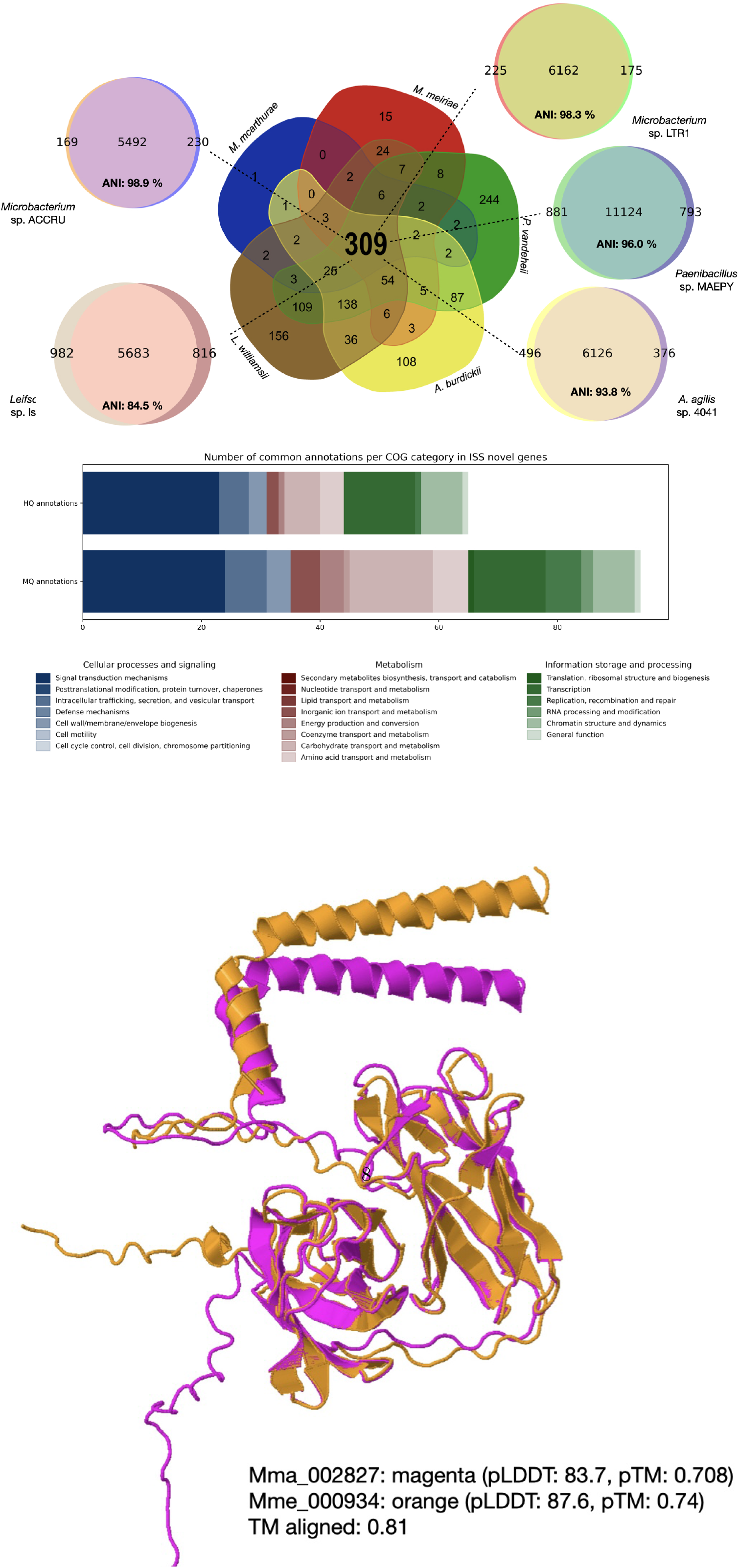

When aligning the novel genes in the genome context, we found the presence of potential novel biosynthetic gene clusters (BGCs) in all species. These BGCs encode secondary metabolites such as terpenes, carotenes, betalactones, lanthipeptides, lassopeptides, non-ribosomal metallophores, furans, non-alpha poly-amino acids (NAPAA), type 3 polyketide synthase (T3PKS), ribosomally synthesized and posttranslationally modified peptides (RiPP), proteusins (Supplementary Fig. S1), Our findings of BGCs are complimentary to the previously reported BGCs in *A. burdicki* i, *L. williamsii* and *P. vandeheii* [28]. We also expand the BGCs classification of the studied ISS species into the “novel” category, which are not found in the genomes of their closest terrestrial relatives. In this study, we report small molecule biosynthetic pathways, metal ion transporters and other transmembrane transporters. Furthermore, transcription factors associated with environmental stress have been found (Supplementary Table S14).

Given the difference in ANI for each ISS-species:closest-relative pair, a different number of novel gens was obtained in each case, proportional to the phylogenetic distance (the lower the ANI, the more novel genes were found). We therefore focused on the two *Microbacteria* species, as their ANI to the closest relative was higher than 98%. Using Muscle, we aligned sequences of novel genes from each species together. We found an example of novel genes (Mma 002827 and Mme 000934) being present in both species with high (52.8%) sequence similarity. Given that initial annotation (“heavy metal associated domain”, “response to copper ion”, see Supplementary Tables S2 and S6), was sequence-based (see Methods for details), another approach was assayed, with structure prediction, followed by a functional annotation. Superpositions of predicted structures (both with pTM*>*0.7 and pLDDT*>*80) are almost identical (Fig. 3c). The structure-based deepFRI annotations reveal that these proteins have oxidoreductase activity (acting on CH-NH2 group of donors) and that are located in the extracellular layer of the cell (Supplementary Table S15). When compared to the proteome of *D. radiodurans*, weak structural homology was found to extracellular protein WP 027479552.1 (TM-score 0.34, see Supplementary Fig. S2).

Mobile genetic elements (MGEs) such as transposases were also found in all species except *L. williamsii*. There is also evidence of plasmid in some of the species (*M. mcarthurae, M. meiriae*), due to the presence of relaxosome encoding genes, such as MobC/L family relaxase (Figure 5, Supplementary Table S16).

### 2.2 Comparison of proteome-level mutations in the ISS species and within pangenome context

Several groups of proteins, important for adaptation to space environmental conditions were analyzed between ISS species and their closest Earth relatives. In metallopeptidases, the mutation rate of hydrophobic amino acids and glycine are higher than 1 in this protein group, suggesting an enrichment of these amino acids in the metallopeptidase proteins of ISS bacteria. These mutations might enhance the stability of these proteins under space environmental conditions (Figure 4A). Glycine enrichment is a measure of how often the amino acid glycine appears in a protein sequence compared to what would be expected by chance (see Methods for more details). Significant glycine enrichment in a protein can suggest evolutionary pressures and adaptive responses to make proteins more flexible via mutation to glycine[29]. The metallopeptidase activity stands out as it exhibits consistent glycine enrichment across all strains, suggesting it may play a critical role in universal adaptive mechanisms (Fig. 4B). Other molecular functions, such as “ATP-dependent peptidase activity” and “2-C-methyl-D-erythritol-4-phosphate cytidylyltransferase activity”, show high glycine enrichment, but only in both *Microbaterium* species, indicating potential taxonomy-based adaptations (Fig. 4B).

**Fig. 4.**
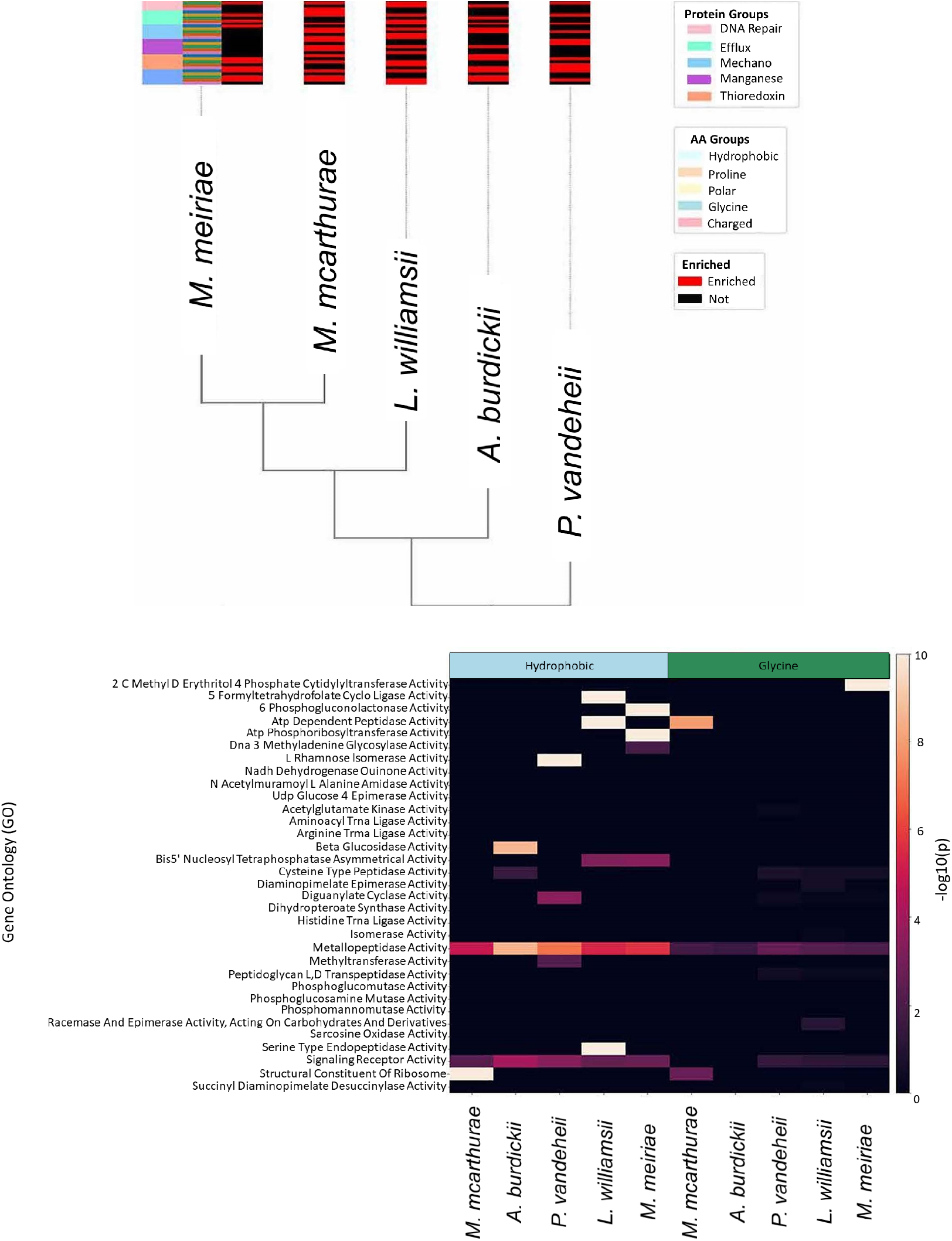
Comparison of mutation enrichment in ISS species across different protein families. A) Clustering dendrogram representing enrichment in protein mutations (wrt closest relative) related to the protein group and type of mutation B) Mutation enrichment of selected proteins (wrt Uniprot database) with their significance scores.

**Fig. 5.**
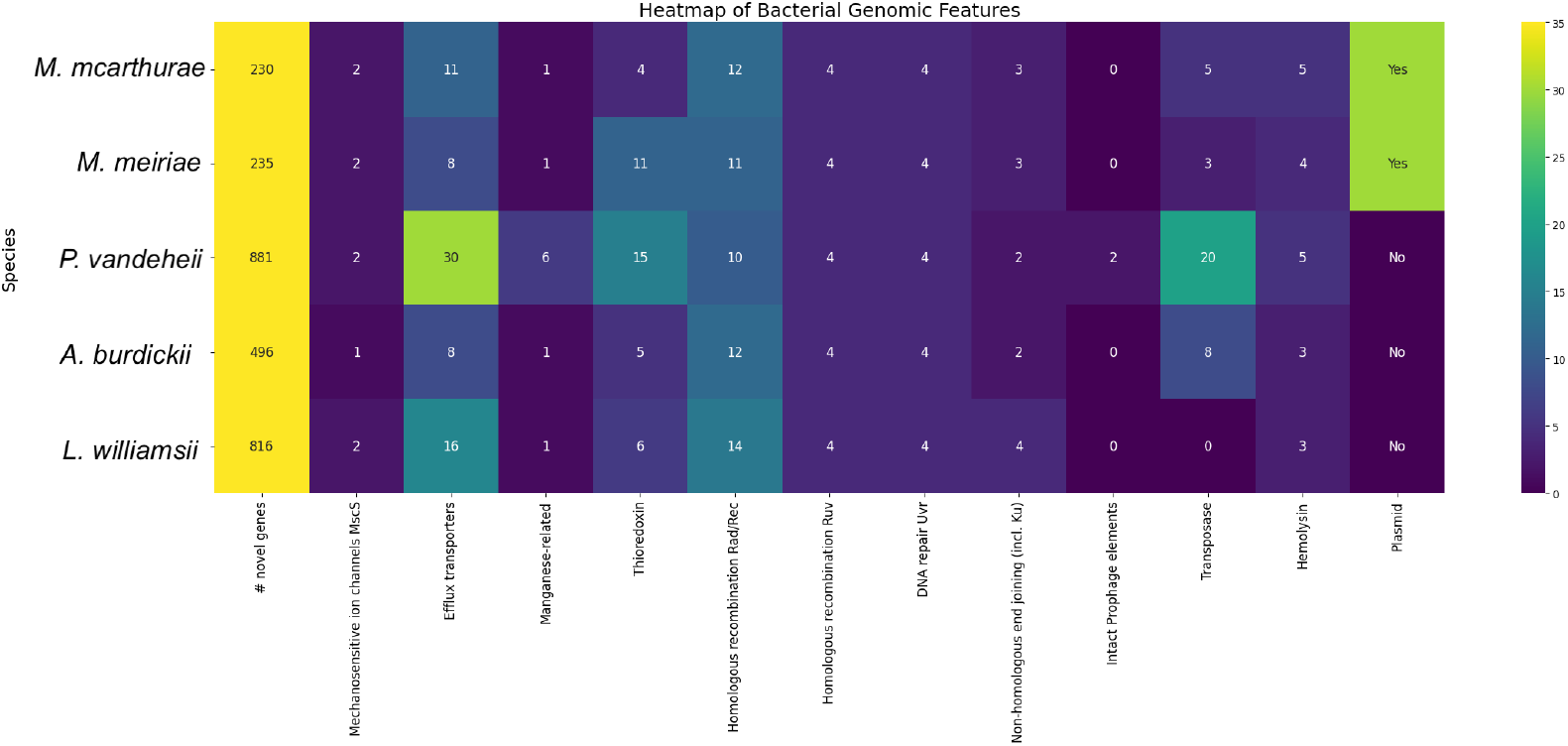
Heatmap representing abundance of annotations of interest in the studied species.

**Fig. 6.**
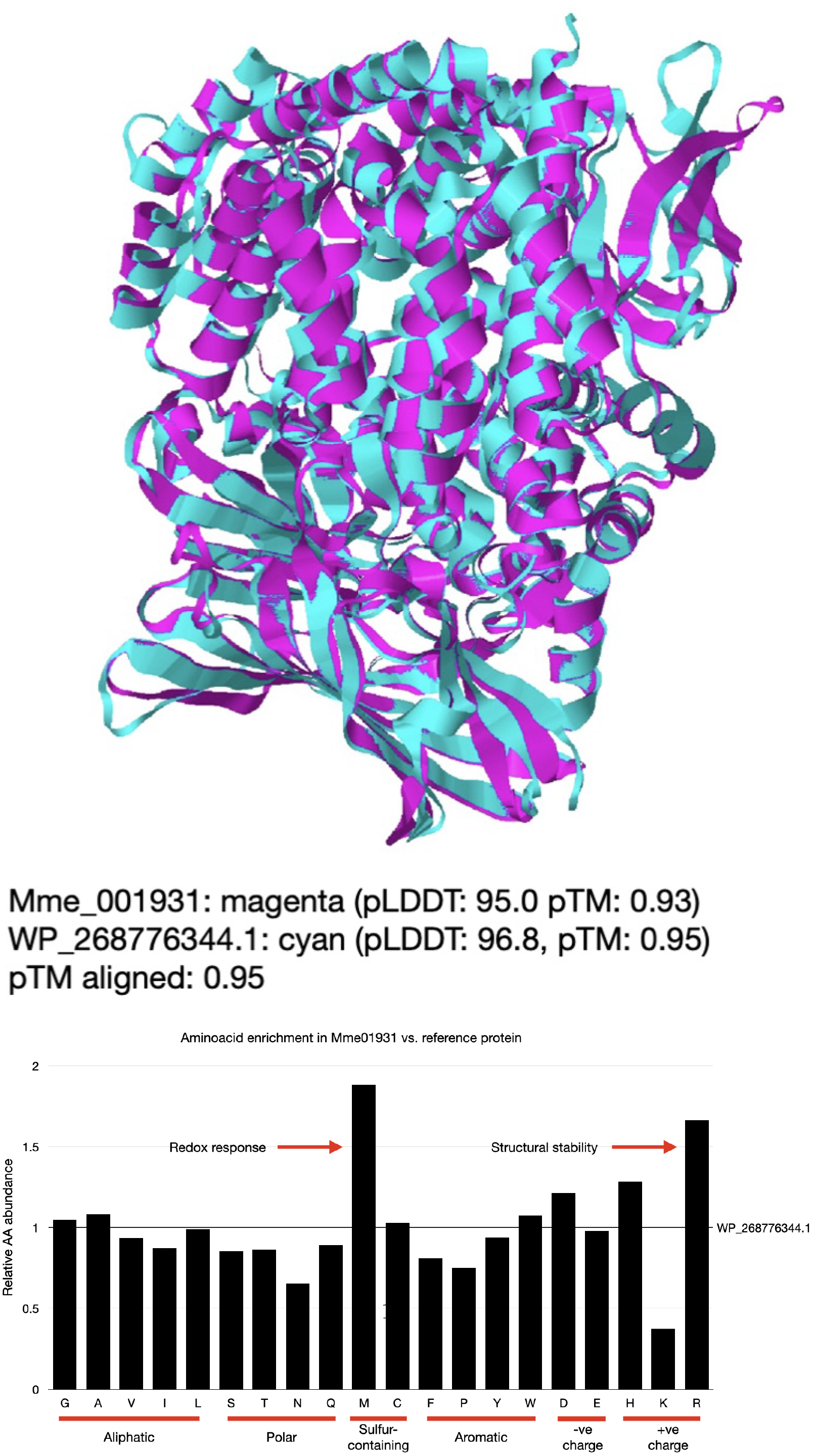
Structure alignment of selected metallopeptidases from *Microbacterium* species, with amino acid enrichment in space isolate. amino acids were grouped by their side chains properties. **pLDDT** - predicted local-distance difference test, **pTM** - predicted template modelling, **TM** - template modelling, see [26, 27]

Certain molecular functions, such as “5 formyltetrahydrofolate cyclo ligase activity”, “ATP dependent peptidase activity”, and “ATP phosphoribosyltransferase activity”, display a high degree of hydrophobic amino acid enrichment, but only in *M. meiriae* and *L. williamsii*. This could suggest convergent adaptations involving changes in hydrophobicity of aforementioned proteins. Interestingly, “metallopeptidase activity” shows hydrophobic amino acid enrichment across all ISS species, albeit at a lower level than glycine enrichment. This could suggest a universal, but less dominant, adaptive strategy involving changes in protein hydrophobicity. These patterns underscore the complexity of potential adaptive strategies, highlighting the interplay between different types of amino acid changes in response to ISS environments (Figure 4B). Metallopeptidases play role in celluar stress response [15], biofilm formation [30], as well as virulence [31] and hence are important factors in microbial adaptations to space conditions. Therefore, a comparative analysis of sequences and structures of selected metallopeptidases from *M. meiriae* and its closest relative will be further discussed in the section below.

### 2.3 Case studies of adaptations

The functional annotations were further investigated to search for genes that would allow adaptation to life in space. The genes of interest were summarized in Figure 5.

#### 2.3.1 Energy metabolism

As illustrated in Fig.1, metabolic rearrangements are an essential part of adaptations to life in space, which was also reported earlier [8–10, 12, 13, 18]. Common annotations show the presence of genes encoding proteins related to carbohydrate, cofactors and coenzymes metabolism (Fig 2c, Fig. 3b). Apart from genomic annotation (Supplementary Table S12), metabolic tests conducted on the ISS bacterial species reveal their ability to grow on a broad range of carbon sources, including cyclic compounds, oligo- and polysaccharides (Supplementary Fig. S3a-b). All species are capable of growing on 27 substrates being the sole carbon source, including carbohydrates (pectin, dextrin, mannose, fucose, gentibiose, lactose, melibiose, glucose, galactose, fructose as well as their branched and glycosylated derivatives), fatty acids (acetic acid, L-malic acid, sodium lactate and butyrate), amino acids (L-glutamic acid), alcohols (glycerol) and antibiotics (vancomycin and aztreonam). They can also grow at 1% NaCl concentration and pH6. Apart from the spore-forming *P. vandeheii*, the other bacteria can also utilize tween 40, gelatin, D-malic acid, L-lactic acid, propionic acid, D/L-serine, L-alanine, sorbitol, lincomycin, minocycline, nalidixic acid, rifamycin, as well as grow in the presence of lithium chloride. On the other hand, only *P. vandeheii* is able to grow in the presence of sodium bromate. Additionally to the species described by Simpson and colleagues [28], two species of *Microbacterium* were characterized for the first time. These species are able to utilize 72 carbon sources altogether (80 and 73 substrates for *M. mcarthurae* and *M. meiriae*, respectively), grow at 8% NaCl concentration, as well as in the presence of lithium chloride and potassium tellurite. Unlike other species, either *M. mcarthurae* or *M. meiriae* can utilize *α*-keto-glutaric acid and citric acid.

#### 2.3.2 Mechanosensitive ion channels (Msc) and osmoprotectants

Except for *A. burdickii*, each of the studied organisms has at least one copy of *msc*S gene annotated. Msc proteins are known to play a crucial role in adapting cells to the osmotic pressure [32]. Overall, two variants of Msc proteins exist: large conductance, (3.6 nS) MscL and small conductance (1 nS) MscS. Large conductance channels are generally conserved and present in most bacteria as well as eukaryotes, although they are not found in deep sea species, where no pressure changes occur. Conversely, MscS are diverse structurally and functionally and occur in those organisms, which exhibit hypo-osmotic stress periodically (e.g. desiccation). MscS proteins have a common core architecture but show different gating mechanisms and conductive properties. Because low gravity conditions can be viewed as decrease in osmotic pressure, leading to the hypo-osmotic stress, their presence is therefore crucial for survival in ISS. All terrestrial relatives have a copy of mscS gene with over 90% sequence similarity to their homologous gene from the ISS species (Supplementary Fig. S4), except for *L. williamsii*, where there is an additional copy of the mscS with no homology in the genome of *Leifsonia* sp. ku-ls. Nevertheless, the presence of an additional copy of mscS gene could be due to the larger phylogenetic distance, as indicated by ANI (Fig. 3A). As for the osmoprotectants, there is no commonly shared metabolite, as genes related to betaine, choline, ectoine and trehalose synthesis are not present in all five species.

#### 2.3.3 Redox stress

Changes in osmotic pressure, as well as exposure to ionizing radiation may lead to the formation of reactive oxygen species (ROS) that have detrimental effects on cells. From the studies on *Deinococcus* it was found that it has elevated intracellular concentrations of Mn as well as Fe[33]. All studied ISS strains show an elevated number of metalloproteins compared to their closest terrestrial counterparts (Supplementary Table S12). In the case of *M. mcarthurae* and *M. meiriae* there are 27 and 25 unique metalloproteins, respectively (Supplementary Table S14), which include metallopeptidases, thioredoxins, metalloregulated transcription factors and heavy metal associated P-type ATPases. Metallopeptidases were of our special interest, as they showed certain adaptive mutations in all studied species (as described in previous section). Therefore, we decided to further analyze the metallopeptidase pair from *M. meiriae* and its terrestrial relative *Microbacterium* sp. LTR1. We found aminopeptidases in two of these species (Mme 001931 and WP 268776344.1 from *M. meiriae* and its terrestrial relative *Microbacterium* sp. LTR1, respectively), which share only 44% protein sequence similarity (58.4% nucleotide similarity, Supplementary Fig. S5). When structural models were calculated for these proteins (both with pTM-score ≫ 0.9), their superposition is almost identical (Fig. 5A), despite the low sequence similarity. When analyzing enrichment of the amino acids in Mme 001931 with WP 268776344.1 as a baseline, we observed enrichment of arginine and methionine, which were over 1.5 times more abundant compared to terrestrial protein. Opposite case was observed for lysine, which was over 2 times less abundant in ISS organism (Fig. 5B). When superimposed to deinococcal aminopeptidase (WP 0516188611), Mme 001931 exhibited similarly high TM-score (0.83), although the latter’s size is 1.81 times bigger than WP 0516188611 (Supplementary Fig. S6). In terms of amino acid enrichment, WP 051618861 also showed an elevated amount of arginines (8.3% wrt 5.5%) with subsequent decrease of lysines (1.1% wrt 2.6%) and increase in methionine with decrease of negatively charged amino acids, but also increase in glycines, histidines and prolines. Nonetheless, there is almost two times the size difference and further phylogenetic distance than in the case of *M. meiriae*.

#### 2.3.4 DNA protection, repair and recombination

Another feature associated with ISS is the elevated rate of DNA-damaging due to radiation. *Deinococcus radiodurans* was extensively tested for its ability to protect and repair damaged DNA, which was due to its homologous recombination (HR) kit. All ISS species encode 10-14 genes related to HR, compared to 10 genes identified in *D. radiodurans* (Fig. 5). Similarly, all of the genomes share the set of *rec, ruv, rad* and *uvr* genes. In terms of non-homologous end-joining (NHEJ), all ISS strains share at least one gene. When compared with the Rec protein family from *D. radiodurans* and *B. pumilus* (Supplementary Fig. S7), clusters reflect phylogenetic distances. Moreover, all ISS strains share the LexA repressor (EC 3.4.21.88), which is part of the SOS response.

#### 2.3.5 Phage elements

Phage elements were found in all genomes of ISS strains, with at least two phage-related genes found in each strain. Holin is a short protein containing few transmembrane regions. Although holin is usually accompanied by endolysin, there is no evidence of phage-related endolysin in any of the genome of ISS strains, suggesting that the phage elements could have been domesticated or removed. When tested with PHASTER software, prophage elements were detected only in the genome of *P. vandeheii*, as well as in its closest Earth relative (Supplementary Table S16).

#### 2.3.6 Potential pathogenicity and antibiotic resistance

Common annotations (Supplementary Table S2) indicate the presence of genes with virulence potential, such as metallophores and hemolysins (including the group III family, which forms oligomeric complexes), which are present in all ISS strains. However, ISS strains often exhibit enhanced expression of these virulence genes due to the unique selective pressures of the space environment, such as microgravity and increased radiation. In contrast, Earth analogous strains show a greater diversity in these genes, driven by varied environmental challenges and human influences like antibiotic use and pollution ([34]. Genomes of all ISS bacterial species contain genes involved in biofilm formation, potentially posing a pathogenic risk to humans under specific conditions, even though these genera and their closest terrestrial relatives are not typically known to be infectious agents. Comparatively, Earth counterparts also form biofilms, but the selective pressures driving this ability differ. On Earth, biofilm formation is often a response to environmental challenges such as antibiotic exposure and surface colonization in diverse habitats. In contrast, ISS bacteria have shown adaptations favoring biofilm formation and surface interactions due to the unique conditions of the space environment, such as microgravity and limited nutrient availability [35]. Additionally, as Mora *et al*. highlighted in their work [36], the genomic and physiological features selected by ISS conditions, though not directly pertinent to human health, exhibit considerable adaptations for biofilm formation and surface interactions. Their research underscores the importance of minimizing local moisture to prevent the formation of potentially harmful biofilms. Metabolic assays indicate that all ISS bacterial species have the ability to degrade at least two antimicrobial compounds (Supplementary Fig. S3ab), showcasing their resilience in a controlled, high-stress environment. These bacteria also harbor various P-type translocating transportases and metallopeptidases within their pool of novel genes (Supplementary Table S12), suggesting advanced mechanisms for survival and resistance. Similarly, Earth analogs have evolved to degrade antimicrobial compounds, especially in environments heavily influenced by human activity, such as hospitals and agricultural sites [31].

## 3 Discussion

The thorough examination of genomes of five novel ISS bacterial species, when compared with their closest terrestrial counterparts, unveiled clear adaptations to the rigors of ISS space environment. Notably, significant shifts in cell physiology and metabolism were seen. The extent to which genes lacked homologs in their terrestrial relatives was found to be associated with their phylogenetic distance. A specific set of genes was discovered to be present in all isolates but absent in their closest terrestrial relatives. This observation is consistent with earlier findings on the ISS-specific evolutionary patterns [37, 38]. Mannose is a known substrate of glycosylation in prokaryotes[39] and thus may play a role in cell wall modifications, cell adhesion and biofilm formation. Moreover, genes related to beta-mannosidase and mannose activities were up-regulated in high salinity media in the brine shrimp *Artemia salina*[40], suggesting that mannose and its derivatives may be accumulated as organic osmolytes.

Clusters associated with small molecule and peptide biosynthesis in fungi were found in abundance, reaffirming earlier studies into molecular responses to microgravity [11]. The pool of novel genes unique to the ISS species (Supplementary Table S13) showed an abundance of genes related to peptide and small molecule synthesis, transmembrane secretion/transport, and mechanosensitive ion channels. This suggests they might have a defense mechanism against hypoosmotic shocks. Furthermore, novel surface oxidoreductases have been identified in *M. mcarthurae* and *M. meiriae* which are not present in their terrestrial analogs. In light of these findings, a deeper investigation using comparative transcriptomics, proteomics, and metabolomics is recommended. To perform meta-omic analysis while cultivating these microorganisms on ISS would be very challenging. Nevertheless, such studies were conducted before, e.g. Tanpopo mission [15], although this study focused on meta-omic analyses post-exposure. Conducting *in vivo* experiments on microogranisms could be performed via simulating some of the conditions experienced in ISS, e.g. increased radiation, vacuum or microgravity. Ground facilities to recreate these conditions exist and has been utilized in such experiments before ([41–43], also reviewed in [10]). It would therefore be possible to compare gene expression profile under these conditions, protein activity (e.g. extracellular metallopeptidases) or even assay secretion of certain metabolites (e.g. chelators, antibiotics) on indicative media.

Efficient DNA repair and its impact on survival in the extreme conditions has long been studied in many organisms [41, 44, 45]. LexA repressor-mediated response is known to be crucial to DNA protection and repair process [46]. Studies on extremophilic bdelloid rotifers highlighted the linkage of efficient DNA repair mechanisms with the presence of high rates of HGT [47, 48]. The detection of MGEs underscores a significant genetic factor hinting at the selective pressures faced by species isolated from the ISS. Interestingly, the quantity of annotated transposases seems to be in proportion to genome size, with *P. vandeheii* genome showcasing the most substantial presence of MGEs. On the other hand, transposable elements were not detected in all ISS species, thus suggesting that TE-mediated genome rearrangements are not a common route for genomic adaptations to space environment. Given that microgravity and hypoosmotic shock may induce dessication stress, the adaptation mechanisms may be similar to dessication-tolerant organisms, e.g. bdelloid rotifers, where TEs are highly diverged [49] and containing unique ORFs, thus indicating a domestication. Hence, it is possible that in the species described in this study the TEs are significantly diverged. Extremophilic lifestyle provides selective pressure on maintaining genome integrity. Therefore, TEs disruptive activity is expected to be quenched, at least in the non-spore forming species. On the other hand, *P. vandeheii* may have a different adaptation system, being a spore-former such as *D. radiodurans*, which is known to contain TEs that are a part of stress-response[50]. Notably, sequences tied to plasmid-specific genes, such as *mob*C and relaxase, were discerned in both *Microbacterium* species, a feature absent in their terrestrial analogs. This emphasizes the potential unique genetic adaptations of these space-borne bacteria.

Building upon established research[33, 47], our analysis of proteomic alterations points to a significant enrichment across various protein classes. A particularly noteworthy observation is the global glycine enrichment, a phenomenon that aligns with findings from earlier studies on *Acinetobacter* sp., a desiccation-tolerant organism[17, 37, 51]. Historically, these organisms have not only displayed enhanced hydrophilicity but have also shown a fortified resistance to ionizing radiation, a stress factor intrinsically tied to redox reactions. The emphasis on these genetic mechanisms holds paramount importance, especially in the context of space-related stress factors. As organisms venture into space, they confront an array of unique and potent stressors, including elevated radiation, microgravity, and vacuum conditions. The genetic adaptability and resilience exhibited by certain organisms, as demonstrated by their proteomic profiles and specific enrichments, provide invaluable insights. These genetic mechanisms might well be the key to understanding how life can persist, and perhaps thrive, in the harsh environs of space. Such knowledge not only expands our understanding of extremophile biology but also informs potential biotechnological applications and space colonization strategies in the future.

When looking at the singular case of metallopeptidase (aminopeptidase N) from *M. meiriae* and its closely-related Earth counterpart *Microbacterium* sp. LTR1, we observed almost perfect structure alignment, despite 43% sequence similarity. The changes in amino acid composition reflect the adaptations to the lifestyle of *M. meiriae*, with enrichment of arginine(s) and methionine(s), with the subsequent reduction of lysine and asparagine. Arginine is known to provide more stability to the structure than lysine [52–54]. Methionine in proteins plays an important role in cellular antioxidant activity, protein structure stabilization, and sequence-independent recognition of protein surfaces. It can also act as a regulatory switch through reversible oxidation and reduction (reviewed in [55]). This could hence explain its presence in the catalytic site (HEXXH motif) of aminopeptidase N.

Drawing from foundational work such as that by Daly et al.[33], metalloproteins stand out as a consistently enriched protein class in strains adapted to space conditions. This enrichment pattern mirrors the remarkable adaptations seen in organisms like *D. radiodurans*, which has gained attention for its extraordinary radiation resistance[41]. The prominence of these metal-dependent enzymes in space-adapted strains is not merely a coincidence. These enzymes play a pivotal role in efficiently managing the redox stress that such microorganisms face in space. Their enrichment suggests an enhanced capability for early stress detection, with these enzymes acting as both environmental sensors and transcription factors. This is a vital adaptation, as space presents a plethora of unique stressors, with redox imbalances being a major concern. The ability of certain strains to bolster their defense mechanisms, as evidenced by the enrichment of specific proteins, reveals the intricate genetic adaptations that enable survival in such hostile settings. These findings not only deepen our grasp of astrobiology but also have profound implications for potential biotechnological applications, hinting at ways to engineer organisms or systems for better resilience in space or other extreme conditions. The extreme ISS environment explains the need for metals scavenging to balance the redox stress e.g. in the form of metalloproteins. The study of metallopeptidases is essential in the context of microbial adaptation to space conditions due to their significant roles in protein degradation, regulation of biological functions, and contribution to microbial virulence and biofilm formation[30]. This research is vital for ensuring the health and safety of astronauts, maintaining the integrity of space missions, and exploring new biotechnological applications. On the other hand, the prevalence of genes such as hemolysins, ferrochelatases, as well as protein glycosylation (mannose), antibiotic resistance and biofilm pathways indicates the risk of potential pathogenicity of the ISS-inhabiting microorganisms. These genes also appear in the genomes of their closest terrestrial relatives, but more studies needed to validate whether microgravity enhance the antibiotic resistance and biofilm production[56]. Conversely, these observations highlight the importance of metal uptake systems, which could lead to the development of antimicrobial agents utilized in the space habitat. Hemolysins are known to be repressed upon high Fe concentrations, as they are tools to scavenge it from the environment. On the other hand, LexA-mediated stress response is known to initiate bacteriocin synthesis [57]. Thiopeptide antibiotic called TP-1161 has been detected in the ISS organism *A. burdickii* [28] and the presence of BGCs encoding potentially novel antimicrobial molecules in the genomes of remaining four ISS species has been confirmed. Therefore, these organisms may serve as a source of potential new antimicrobial agents available upon spaceflight. In summary, the present study underscored the benefits of deep learning-based annotations, here using DeepFRI[58], which are able to go beyond the limitations of homology-based tools like PGAP and eggNOG[59]. In fact, DeepFRI enabled the construction of a comprehensive and cohesive picture of adaptations to space conditions. Nevertheless, combined use of annotations derived from all tools was applied to e.g. mechanosensitive ion channel proteins (see Section 2.3.2). While traditional tools often relegated these genes to the ambiguous “hypothetical” category, Deep-FRI was adept at providing precise annotations. This allowed for the discernment of novel BGCs, a finding that was further corroborated using specialized tools such as antiSMASH[28, 60]. In addition, a significant portion of these novel annotations pertained to membrane-associated proteins. This resonates with prior observations highlighting membrane-related adjustments. The challenge of studying the membrane proteome cannot be understated. Yet, with the recent augmentation of databases like AlphaFold[27] and ESM atlas[61], the scientific community has been endowed with a richer repository of membrane-bound protein structures. This surge in data availability paves the way for more refined structure-based annotations of novel genomes.

This deep dive into the genetic underpinnings bears profound significance, especially when contextualized within space-related stress factors. As organisms confront the multifaceted challenges of space, their genetic machinery undergoes intricate recalibrations. By harnessing sophisticated annotation tools, we gain unparalleled insights into these genetic adaptations, potentially unlocking the secrets of space resilience. Such revelations not only propel our understanding of astrobiology forward but also spotlight the potential of genetic mechanisms in tailoring organisms for optimal performance in extraterrestrial environments.

## 4 Methods

### 4.1 Bacteria isolation, genome assembly and annotation

Sample collection and bacterial isolation from ISS were performed as described in [23, 62]. Biochemical tests were performed using a Gram-positive identification card (Vitek 2 GP ID, bioMèrieux) and phenotypic fingerprint was generated through GNIII MicroPlate according to BioLog’s protocol as described elsewhere [28]. We assembled the genomes of five novel bacterial species isolated from various surfaces of the ISS (specifically strains IF8SW-P5, IIF3SC B10, F6 8S P 1B, F6 8S P 2B, F6 3S P 1C), using the SPAdes workflow[63]. Phylogenetic affiliations are as outlined in [28] and presented in Table 1.

### 4.2 DNA extraction

The ISS isolates (n=5) were grown on R2A medium and incubated at 30°C, for 2 to 7 days. For WGS, DNA was isolated with the ZymoBIOMICS DNA MagBead kit, and WGS libraries prepared using the Illumina Nextera DNA Flex kit. Sequencing on the NovaSeq 6000 platform involved paired-end, 2 x 150 bp reads. FastQC (https://github.com/s-andrews/FastQC) and fastp [64] ensured data quality. WGS reads were assembled using SPAdes [63], with genome statistics from QUAST [65] and quality checks from CheckM [66]. Species verification through fastANI [67] and fastAAI (https://github.com/cruizperez/FastAAI) confirmed identities with over 95% identity, and digital DNA-DNA hybridization (dDDH) was estimated with GGDC using Formula 2[68]. For genome synteny analysis, we used SynTracker [25].

### 4.3 Genome annotation

Following assembly, the contigs were subjected to the NCBI Prokaryotic Genome Annotation Pipeline (PGAP)[69], available at https://github.com/ncbi/pgap/wiki, in order to obtain draft genomes and proteomes. Genomes were further investigated for their functions, where all identified genes (bearing PGAP descriptions in their names) underwent homology-based annotation using eggNOG v5.0, accessible at https://github.com/eggnogdb/eggnog-mapper. Genomes of closest relatives of the ISS species (highest average nucleotide identity [ANI]) were also subjected to PGAP and eggNOG annotation.

Proteomes with the PGAP identifiers were used as input for DeepFRI analysis [58], available on github: (https://github.com/bioinf-mcb/Metagenomic-DeepFRI/), which generated query contact map using results from mmseqs2 target database[70] search for similar protein sequences with known structures and with v 1.0 model weights. For reference database, we combined structures from Swissprot (https://alphafold.ebi.ac.uk/download), microbiome immunity project (MIP) [71] and protein databank (PDB, https://www.rcsb.org). Next, the contact map alignment was performed to use it as input to DeepFRI’s Graph Convolutional Network (GCN) and annotations without the alignment to known structures were processed with Convolutional Neural Network (CNN). For medium quality (MQ) and high quality (HQ) annotation, scores higher than or equal to 0.3 (50% of prediction confidence; MQ) and 0.5 (90% of prediction confidence; HQ) were used respectively (Figs. 2-3). All annotations for ISS strains and their terrestrial relatives are provided in Supplementary Tables S2-S11. Phylogenetically close relatives from Earth for each tested species were identified using ANI (Table 1). Their genomes were annotated using DeepFRI as described above. Protein sequences from the aforementioned five species were aligned and conducted two-dimensional clustering using CD-HIT[72], available at https://github.com/weizhongli/cdhit, employing a 40% similarity threshold. Genes below that threshold were considered “novel” and subjected to analysis shown in Fig.3A-B.

Annotations (GO terms) were then compared between ISS strains (Supplementary Table S12) and their terrestrial counterparts (Supplementary Table S13). Common annotations were aggregated according to clusters of orthologous genes (COG) categories[73] using go2cog mapper (https://github.com/Tomasz-Lab/go2cog). GO terms that could be assigned to more than 3 COG categories were grouped as category R - “general function prediction” (Figures 2 and 3). Heatmaps were generated using seaborn (https://github.com/mwaskom/seaborn). Venn diagrams and Upset-plot were generated using matplotlib (https://github.com/matplotlib/matplotlib). Using a knowledge-based approach, a set of genes was selected for further analysis. Protein structures were modeled using *in silico* structure prediction tool – AlphaFold Colab[27] – and hence obtained protein databank (PDB) format structures were aligned using Geneious Prime 2023.1.1 (Biomatters). Structures were aligned using TM-align tool [26]. Based on the predicted structure, novel functional annotation was performed using DeepFRI. All the annotation data and scripts are placed in the repository on github (https://github.com/lmszydlowski/Adaptation-of-novel-bacterial-species-isolated-from-the-International-Space-Station).

### 4.4 Sequence alignments, functional annotations for plasmids, phage elements and biosynthetic gene clusters in the genome

Sequences of selected proteins were aligned using Muscle 5.1 [74]. Clustering dendrograms were built using PhyML 3.3,[75] using LG amino acid substitution model and approximate likelihood ratio test (aLRT) for branch support. All contigs generated for each genome were scanned for potential plasmids using PLASme software (https://github.com/HubertTang/PLASMe), phage-related elements using PHASTER software[76, 77] and results provided in the Supplementary Tables S15-S16. Genome sequences were also processed through the AntiSMASH 7.0 software [60] to look for biosynthetic gene clusters (BGCs) (Supplementary Table S14).

### 4.5 Protein mutation and enrichment analysis

To investigate the roles and processes of proteins highly enriched in bacteria from the ISS, we specifically focused on the enrichment of certain amino acids, especially proline and glycine, which belong to unique amino acid change groups. The amino acid compositions of proteins from the ISS strains were compared with those proteins found in the closest terrestrial species through global alignment methods[78]. This facilitates the identification and quantification of any shifts in the amino acid composition. These shifts were evaluated by calculating the mutation rate, which is achieved by dividing the number of amino acid substitutions in the ISS bacteria by the number of amino acid substitutions in their closest terrestrial relatives. Proteins amino acid mutation rate normalized and Z-score above 1 .96 or below -1.96 were classified as enriched. To investigate the functions and pathways of these proteins, we utilize the g:Profiler enrichment tool (https://biit.cs.ut.ee/gprofiler/gost[79]). Custom gene sets for bacteria were created using UniProt’s[80] predicted Gene Ontology (GO) term associations[81] for bacterial proteins. The tool was configured to include a minimum gene set size of 5 and a maximum of 10,000, focusing on molecular function GO terms. Shared gene sets between samples were filtered using a false discovery rate (FDR) cutoff of 0.05 in 1424 molecular function GO terms. Data visualization was enhanced through ete3 (https://doi.org/10.1093/molbev/msw046) for constructing phylogenetic trees and seaborn (https://github.com/mwaskom/seaborn) for generating heatmaps. The phylogenetic distance between bacterial species was calculated using NCBI’s distance calculations, with the closest terrestrial bacteria as a reference.

## Supplementary information

Supplementary information contains 7 figures and 16 Tables.

**Supplementary Figure 1**. Assembled contigs with regions corresponding to biosynthetic gene clusters marked. Full description of BGC from each organism is provided in the Supplementary Table S14.

**Supplementary Figure 2**. Alignment of novel S-layer osidoreductases with their closest sequenctial homolog from *D. radiodurans* (WP 027479552.1, red). Mma - *M. mcarthurae* (magenta), Mme - *M. meiriae*, orange. **pLDDT** - predicted local-distance difference test, **pTM** - predicted template modelling, **TM** - template modelling, see [26, 27].

**Supplementary Figure 3. A)** Phenotypic assays of studied organisms with GNIII MicroPlate (Biolog). **B)** Biochemical assays of studied organisms Gram-positive identification card Vitek 2 GP ID (bioMèrieux). Blue - absent, red - present. Assays were performed according to manufacturers’ protocol.

**Supplementary Figure 4**. Phylogenetic tree based on alignment of MscS protein sequences from ISS organisms and their closest terrestrial relatives. Branch labels indicate substitutions per site. Ab - *A. burdickii* (blue), Asp - *A. agilis* 4041 (blue), Mma - *M. mcarthurae* (burgundy), Msp - *Microbacterium* sp. ACCRU (burgundy), Lw - *L. williamsii* (yellow), Lsp - *Leifsonia* sp.ku-ls (yellow), Mme - *M. meiriae* (orange), Mp - *Microbacterium* sp. LTR1 (orange), Pv - *P. vandeheii*, (green), Psp - *Paenibacillus* sp. MAEPY (green).

**Supplementary Figure 5**. Alignment of DNA and amino acid sequence of aminopeptidases Mme 001931 and WP 268776344. Green boxes indicate sequence identity.

**Supplementary Figure 6**. Structure alignment of aminopeptidase Mme 001931 (cyan) and WP 0516188611 (red). **pLDDT** - predicted local-distance difference test, **pTM** - predicted template modelling, **TM** - template modelling, see [26, 27].

**Supplementary Figure 7**. Phylogenetic tree based on alignment of DNA-repair protein sequences from ISS organisms and *D. radiodurans* and *B. pumilus* SAFR-032. Branch labels indicate substitutions per site. Ab - *A. burdickii* (blue), Bp - *B*.*pumilus* (black), Dr - *D. radiodurans* (red), Mma - *M. mcarthurae* (burgundy), Lw - *L. williamsii* (yellow), Mme - *M. meiriae* (orange), Pv - *P. vandeheii*, (green).

**Supplementary Table 1**. ANI results (top 8) for all ISS species.

**Supplementary Table 2**. Annotations of *M. mcarthurae* genes.

**Supplementary Table 3**. Annotations of *A. burdickii* genes.

**Supplementary Table 4**. Annotations of *P. vandeheii* genes.

**Supplementary Table 5**. Annotations of *L. williamsii* genes.

**Supplementary Table 6**. Annotations of *M. meiriae* genes.

**Supplementary Table 7**. Annotations of *Microbacterium* sp. ACCRU genes.

**Supplementary Table 8**. Annotations of *A. agilis* 4041 genes.

**Supplementary Table 9**. Annotations of *Paenibacillus* sp. MAEPY2 genes.

**Supplementary Table 10**. Annotations of *Leifsonia* sp. ls-ku genes.

**Supplementary Table 11**. Annotations of *Microbacterium* sp. LTR1 genes.

**Supplementary Table 12**. Annotations of genes common in all ISS species.

**Supplementary Table 13**. Annotations of unique (not present in closest relative) genes shared by all ISS species.

**Supplementary Table 14**. DeepFRI annotations of unique (not present in closest relative) genes and antiSMASH predictions of BGCs.

**Supplementary Table 15**. Structure - based DeepFRI annotations of Mma 002827 and Mme 000934.

**Supplementary Table 16**. Predictions for plasmids and phage elements in the genomes of ISS species.

## Acknowledgements

We acknowledge the Jet Propulsion Laboratory supercomputing facility staff, notably Narendra J. Patel (Jimmy) and Edward Villanueva, for their continuous support in providing the best possible infrastructure for BIGDATA analysis. © 2023 California Institute of Technology, Government sponsorship acknowledged.

## Declarations

### Ethics approval

Not applicable

### Consent to participate

Not applicable

### Consent for publication

All authors have agreed to be co-authors and have approved the final version of this manuscript.

### Availability of data and materials

The draft genome sequences of strains IIF3SC B10, F6 3S P 1C and F6 8S P 1B were deposited in NCBI under BioProject PRJNA935338 (see [28]. Strains F6 8S P 2B and IF8SW-P5 were deposited in BioProjects PRJNA869446 and PRJNA224116, respectively. The WGS accession numbers are given in Table 1 and the genome versions described in this paper are the first versions. Moreover, genomes of all ISS strains and their terrestrial relatives are available on Kbase (narrative 107942).

### Code availability

All bioinformatic tools used in this study are available on GitHub: https://github.com/lmszydlowski/Adaptation-of-novel-bacterial-species-isolated-from-the-International-Space-Station.

### Authors’ contributions

- **LS**, manuscript writing, data analysis, figure preparation;
- **AB**, figure preparation, data analysis;
- **AS**, data analysis, manuscript reviewing;
- **NS**, data analysis, manuscript reviewing;
- **DEK**, figure preparation, manuscript reviewing;
- **PL**, manuscript reviewing, project supervision;
- **OUS**, manuscript reviewing, project supervision;
- **TK**, manuscript reviewing, data analysis, project supervision;
- **VK**, conceptualization, manuscript reviewing, project supervision;

